# SLFN11 restrains alternative lengthening of telomeres

**DOI:** 10.64898/2025.12.08.692725

**Authors:** Prashant Khandagale, Allison L. Welter, Sarah F Clatterbuck, Shar-yin Naomi Huang, Daiki Taniyama, Suresh Kumar, Yuichi J. Machida, Yves Pommier

## Abstract

Alternative lengthening of telomeres (ALT) is a homologous recombination-dependent mechanism maintaining telomere length in approximately 10-15% of all cancers that are telomerase (TERT) negative. ALT is most prominent in osteosarcoma. Although many ALT cells feature loss of the ATRX/DAXX chromatin remodeling complex, ATRX/DAXX deficiency alone is insufficient to trigger ALT. Here, we provide evidence that Schlafen 11 (SLFN11) acts as a suppressor of the ALT pathway. TERT-negative osteosarcoma U2-OS (ALT) cells that normally lack SLFN11 expression show SLFN11 localization to telomeres upon doxycycline-induced SLFN11 expression. This re-expression suppresses ALT activity, as evidenced by reduced ALT-associated PML bodies (APBs), c-circles and telomere DNA synthesis along with decreased levels of Telomeric Repeat-containing RNA (TERRA). SLFN11 re-expression also attenuates the telomeric DNA damage response (DDR) and induces telomere destabilization in ALT cells. Furthermore, SLFN11 suppresses ALT induction in ATRX-depleted prostate carcinoma DU145 and EW8 cells. Collectively, our findings identify SLFN11 as a negative regulator of the ALT pathway. These findings suggest that lack of SLFN11 expression together with ATRX/DAXX inactivation contributes to ALT activation, and that SLFN11 reactivation may be considered to eliminate ALT-dependent cancer cells.

## Introduction

Telomeres are the specialized nucleoprotein structures localized at the distal ends of eukaryotic chromosomes. They are composed of tandem repeats of the hexanucleotide sequence 5’-TTAGGG covered by an assemblage of proteins called the shelterin complex ^1^. The primary function of telomeres is to preserve genomic integrity by preventing chromosomal ends from being shortened during DNA replication and misrecognized as DNA double-stranded breaks. In the absence of this protection, the cellular DNA damage response (DDR) machinery inappropriately engages at chromosomal ends, leading to end-to-end chromosomal fusions ^2^.

In normal somatic cells, the telomeres undergo continuous trimming with each cell cycle due to the incomplete replication of chromosomal ends. When the telomeres shorten to a critical threshold and cells attempt to replicate, cells undergo senescence followed by death. In cancers, telomere elongation is crucial to achieve replicative immortality. While most cancer cells reactivate telomerase (TERT) to maintain telomere length, certain cancers such as sarcomas, glioblastomas and adenocarcinomas commonly lack telomerase and instead utilize a break-induced homologous recombination (BIR)-based mechanism known as alternative lengthening of telomeres (ALT) ^3–9^. Telomere elongation in ALT cells is dependent on the telomeric recruitment of recombination/replication factors including RPA, BLM, TOP3A, RMI1 and RMI2 (BTRR) ^10–14^. Typical features of ALT cells, which differentiate them from telomerase-positive cancers and somatic cells, include accumulation of partially double-stranded circular DNA (c-circles), enriched levels of telomeric RNA (TERRA) at telomeres, formation of ALT-associated PML bodies, telomere heterogeneity, and spontaneous DNA damage signaling with accumulation of phosphorylated histone H2AX (γH2AX) at telomeres ^15–19^.

The mechanisms by which cancer cells activate the ALT pathway remain incompletely understood. One of the major insights into ALT activation is that not all but a significant fraction of ALT-positive cancers are associated with the loss of the chromatin remodeling factor ATRX and/or of the histone chaperone protein DAXX ^19–21^. Together ATRX and DAXX form a complex that deposits the histone variant H3.3 at heterochromatic DNA, including the sub-telomeric and telomeric regions ^22–24^. Although re-expression of ATRX protein in ATRX-deficient ALT cancer cells suppress the ALT phenotypes^25^, the loss of ATRX/DAXX in telomerase positive cells is insufficient to activate the ALT pathway, underscoring that additional, yet unidentified, factors are required for ALT activation ^19^.

Spontaneous DNA breaks at telomeres can also trigger ALT. Secondary structures such as G-quadruplex DNA, telomeric D-loops and TERRA R-loops cause replication fork collapses that provide primary substrates for break-induced telomere elongation ^26–28^. The accumulation of damaged telomeres in PML nuclear bodies provides a recombinogenic microenvironment for homologous recombination, ultimately driving telomere amplification ^29^. Accordingly, induction of telomere breaks by TRF1-FOK1 activates ALT through an accumulation of TERRA R-loops at telomeres ^29,30^. Telomeric DNA stress either through DNA protein-links-inducing agents or oxidative damage stimulates ALT in cells lacking ATRX ^31,32^. These observations suggest that induction of telomeric DNA damage together with inactivation of the ATRX-DAXX complex represent a key requirement for the activation of ALT.

Because factors that suppress ALT are not fully understood, we explored whether Schlafen 11 (SLFN11), which binds single-stranded DNA (ssDNA), RPA (Replication Protein A) and RNA and blocks stressed replicons ^33–35^, plays a role in suppressing ALT. SLFN11, a member of the interferon-inducible Schlafen family, is epigenetically silenced in approximately 50% of human cancer cell lines and cancer tissues ^36,37^. Its expression is causally associated with sensitivity to a wide range of DNA-damaging chemotherapies targeting DNA replication, including inhibitors of ribonucleotide reductase (hydroxyurea), inhibitors of thymidylate synthetase and chain elongation (cytarabine, gemcitabine), PARP inhibitors (talazoparib, olaparib, niraparib), topoisomerase inhibitors (topotecan, exatecan, irinotecan, etoposide, doxorubicin) and platinum compounds (carboplatin, cisplatin) ^33,36,38,39^. Mechanistically SLFN11 possesses an endoribonuclease activity ^34,40^ that selectively degrades the tRNAs required for the translation of critical DNA repair protein kinases such as ATR and ATM ^41,42^. SLFN11 endoribonuclease activity has also been implicated in the regulation of single-stranded nucleic acids including various RNAs ^43–45^. In addition, SLFN11 possesses an ATPase/helicase and single-stranded DNA binding activity that directly inhibits abnormal replication and DNA homologous recombination-mediated repair ^33–35,46^. Because ALT cancers rely on homologous recombination with RPA as a crucial protein involved in the process, we speculated that SLFN11 might control telomeric recombination in ALT cancers ^47^.

## Results

### SLFN11 association with ALT telomeres

To probe the expression of SLFN11 in telomerase-negative cells, which are presumably ALT cells, first we examined the correlations between the expression of SLFN11 RNA and the expression of the telomerase reverse transcriptase (TERT) gene in the publicly available collection of sarcoma cell lines, which are known to be frequently ALT. Using the CellMiner portal (https://discover.nci.nih.gov/cellminercdb/) ^48^, we found that the SLFN11 expression is positively correlated with TERT expression (Pearson coefficient p = 0.001) (Figure 1A), implying that SLFN11 expression tends to be suppressed in ALT cancer cells. Moreover, analysis of the methylome of U2OS, a classical ALT cancer cell line revealed that lack of SLFN11 expression is due to promoter hypermethylation ^38,49,50^ (Figure S1A).

**Figure 1:**
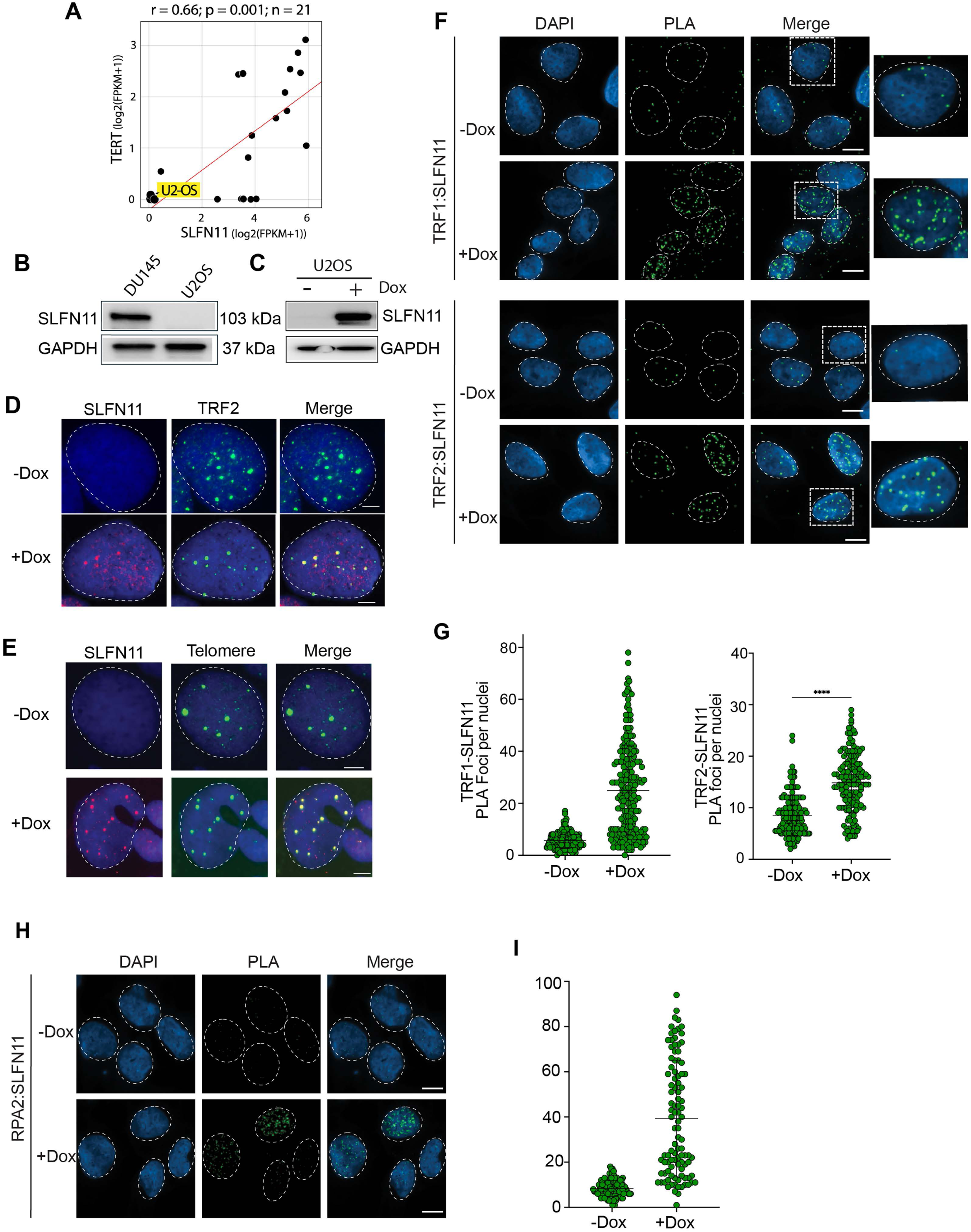
SLFN11 localization at ALT telomeres. **A.** Snapshot image of Sarcoma CellMinerCDB (https://discover.nci.nih.gov/sarcomaCellMinerCDB/) showing that lack of telomerase (TERT) expression (representing ALT cells) is correlated with low and no SLFN11 expression. Figure S1 shows that lack of SLFN11 expression in U2OS is due to promoter methylation. **B.** Representative Western blot demonstrating lack of expression of SLFN11 in U2OS cells. Prostate DU145 cells are used as positive control. **C.** Representative Western blot showing inducible expression of SLFN11 in U2OS cells treated with doxycycline (Dox) for 4 days. **D.** Representative immunofluorescence images showing localization of SLFN11 with TRF2 in U2OS cells after doxycycline induction of SLFN11 for 48 hours. Nuclei are highlighted with DAPI in blue and nuclear periphery with dotted lines. The scale bar represents 5 μm. **E.** Representative immunofluorescence combined DNA-FISH (IF-FISH) image showing localization of SLFN11 with telomeres in U2OS cells after doxycycline treatment for 48 hours. The scale bar represents 5 μm. **F.** Representative image of Proximity ligation assay (PLA) showing interaction of SLFN11 with TRF1 and TRF2 in U2OS cells after doxycycline induction for 48 hours. The scale bar represents 10 μm. **G.** Quantification of TRF1-SLFN11 (n=1) and TRF2-SLFN11 (n=2) PLA foci from one and two independent experiment, respectively. Each green dot represents an individual nucleus. Data are presented as mean ± SDs. ****p < 0.0001 (two tailed pair t-test). **H.** Representative image of Proximity ligation assay (PLA) showing interaction of SLFN11 with RPA2 in U2OS cells after doxycycline induction for 48 hours. The scale bar represents 10 μm. **I.** Quantification of RPA2-SLFN11 (n=1) PLA foci from one experiment. Each green dot represents an individual nucleus.

To study the implication of SLFN11 in ALT, we developed a doxycycline-inducible system to express SLFN11 in U2OS cells. Consistent with the RNA-seq data, U2OS do not express SLFN11 protein under basal conditions (Figure 1B). Upon doxycycline induction, SLFN11 was robustly expressed at levels comparable to those observed in DU145 cells (Figures 1C and S1B). Immunofluorescence imaging showed that SLFN11 is localized to the nucleus upon doxycycline treatment. In most cells SLFN11 appeared as a diffuse nuclear signal. However, a subset of cells (approximately 5% cells) displayed distinct SLFN11 foci that colocalized with both TRF2 and telomeres (Figures 1D, E and S1C). These observations suggested that expressed SLFN11 associates with ALT telomeres.

To confirm the localization of SLFN11 at telomeres, we performed proximity ligation assay (PLA) with the telomere-binding proteins TRF1 and TRF2. SLFN11 showed PLA foci with both TRF1 and TRF2 (Figures 1F and G). As expected ^33,34^, SLFN11 also showed PLA foci with RPA2, validating the assay as a positive control (Figures 1H and I). Together, these results demonstrate that upon forced expression, SLFN11 colocalizes with telomeres in ALT U2OS cells.

### SLFN11 suppresses ALT by inhibiting the recruitment of RPA, BLM and PML bodies to telomeres

In response to DNA damage SLFN11 is recruited to sites of damage through interaction with ssDNA and RPA ^33,34,46^. At ALT telomeres, RPA limits the accumulation of extrachromosomal single-stranded telomeric DNA, probably by blocking nucleases at chromosomal ends ^47^. We found that the SLFN11 colocalizes at telomeres where it tends to suppress the accumulation of RPA (Figures 2A, B and S2A). These results confirm the PLA data presented in Figure 1.

**Figure 2:**
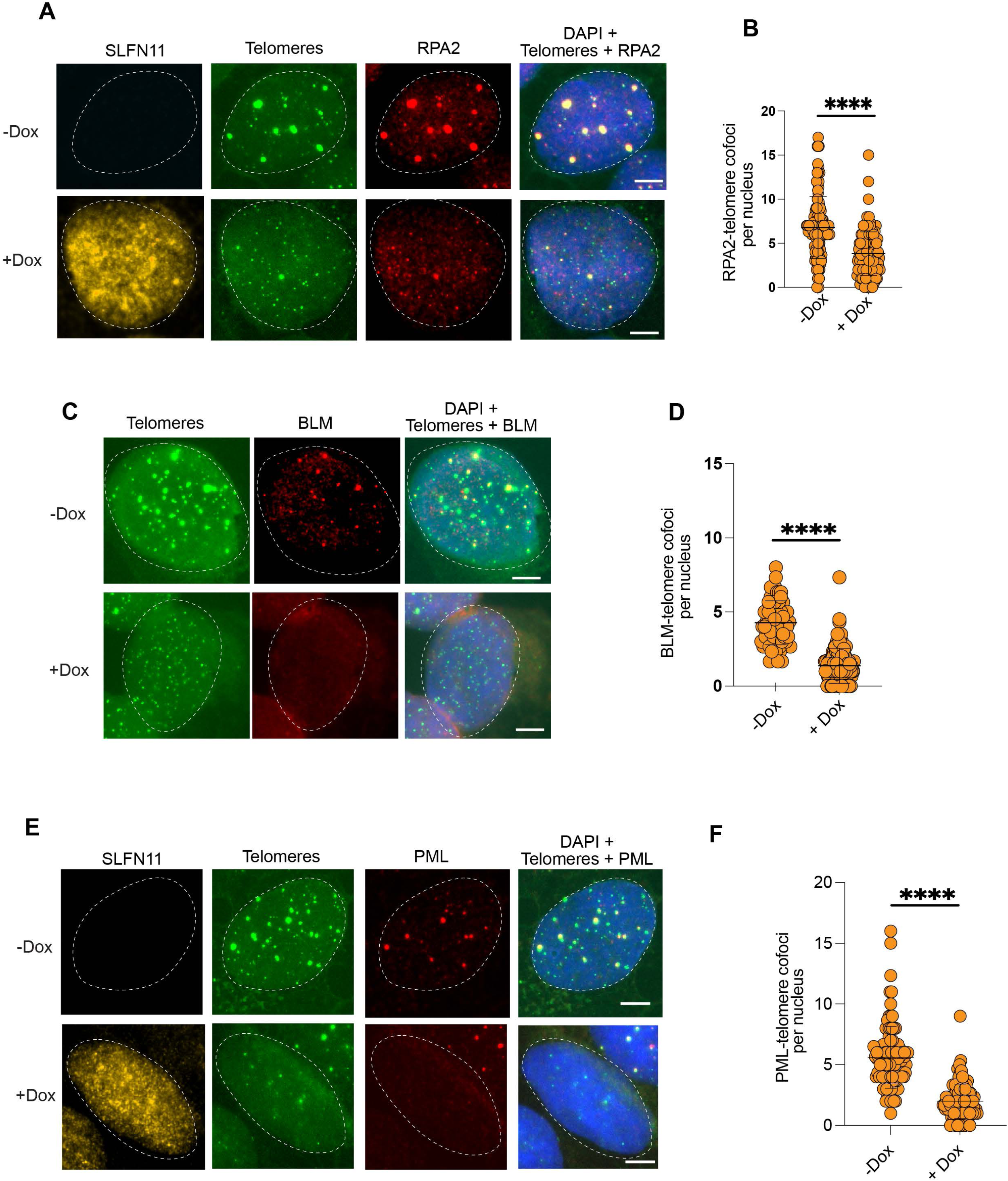
SLFN11 expression impairs the localizations of RPA2, BLM and PML at ALT telomeres. **A.** Representative immunofluorescence combined DNA-FISH images showing RPA2 localization at telomeres without or with SLFN11 expression after 4 days doxycycline treatment of U2OS cells. Scale bar, 5 μm. **B.** Quantification for RPA2 foci at telomeres in U2OS cells with and without SLFN11 expression after 4 days of doxycycline treatment. At least 50 nuclei were counted for each condition in three independent biological experiments. Each orange dot represents an individual nucleus. Data are presented as mean ± SD. **** indicates *p* < 0.0001 (two tailed pair t-test). **C.** Same as panel A with BLM staining. Scale bar, 5 μm. **D.** Quantification for BLM foci at telomeres in U2OS cells with and without SLFN11 expression after 4 days doxycycline treatment. At least 60 nuclei were counted for each condition in three independent experiments. Each orange dot represents an individual nucleus. Data are presented as mean ± SD. **** indicates *p* < 0.0001. (two tailed pair t-test) **E.** Same as panel A for PML. Scale bar, 5 μm. **F.** Quantification for PML foci colocalized at telomeres in U2OS cells with and without SLFN11 expression after 4 days of doxycycline treatment. At least 50 nuclei were counted for each condition in three independent experiments. Each orange dot represents an individual nucleus. Data are presented as mean ± SD. **** indicates *p* < 0.0001 (two tailed pair t-test).

The Bloom helicase is indispensable for ALT telomere elongation, where it not only resolves the recombination intermediates to complete telomere synthesis, but also stabilizes the shelterin complex and promotes Telomeric Repeat-containing RNA (TERRA) R-loops and unwinding ^12,51,52^. At ALT telomeres, a recent study concluded that BLM generates 5′-flaps on the lagging strand, thereby facilitating homologous recombination while protecting telomeric DNA from DNA2 nuclease activity ^52^. Upon SLFN11 expression, we found that SLFN11 suppresses the recruitment of BLM to ALT telomeres (Figures 2C and D).

Another defining feature of ALT is the assembly of SUMO-dependent PML nuclear bodies formed through liquid-liquid phase separation. Those PML bodies serve as specialized compartments where homologous recombination is executed to promote telomere elongation ^53^. We found a marked reduction of the PML signals at ALT telomeres upon SLFN11 expression implying dysfunctional telomere-associated PML bodies in the presence of SLFN11 (Figures 2E, F and S2B).

To exclude the possibility that this reduction was caused by doxycycline itself, we treated parental U2OS cells with doxycycline under the same experimental conditions. Doxycycline treatment did not significantly alter APB formation in parental U2OS cells (Figure S2C), indicating that the reduction in APBs observed following induction is attributable to SLFN11 expression rather than to doxycycline treatment. Together, these results demonstrate that SLFN11 suppresses the recruitment of key ALT-associated factors, including RPA, BLM, and PML bodies, to telomeres.

### SLFN11 decreases the Telomeric RNA and extrachromosomal DNA

In ALT cells, telomeres transcription by RNA polymerase II (RNAP) generates Telomeric Repeat-containing RNA (TERRA) foci consisting of long-non-coding RNA molecules interacting with TRF2 and telomeric R-loops ^12,54,55^, which have been proposed to act as substrates for BIR at telomeres ^56^. The RNA-FISH experiment demonstrates that expression of SLFN11 decreases TERRA foci in U2OS cells (Figures 3A and B).

**Figure 3:**
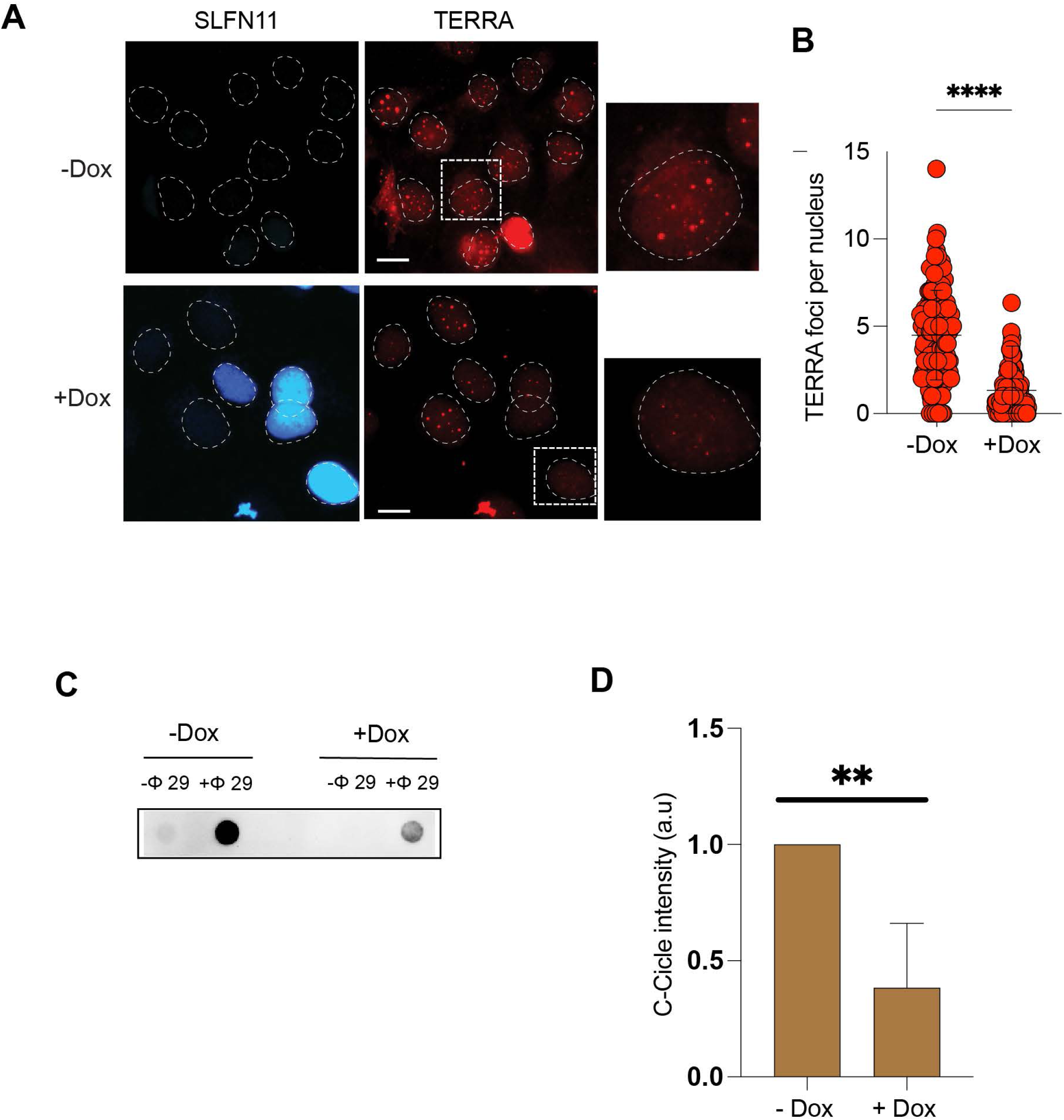
SLFN11 inhibits TERRA and C-circles in ALT cells. **A.** Representative immunofluorescence image showing TERRA signals identified by IF-RNA FISH in U2OS cells expressing SLFN11 after 4 days doxycycline treatment. Scale bar, 10 μm. **B.** Quantification of TERRA foci in U2OS cells expressing SLFN11 after 4 days doxycycline treatment. At least 100 nuclei were counted for each condition in three independent experiments. Data are presented as mean ± SD. **** indicates *p* < 0.0001 (two tailed pair t-test). **C.** Representative slot blot images showing C-circle signals in SLFN11-inducible cells with or without doxycycline induction and in the absence or presence of phi29 DNA polymerase. **D.** Quantification of C-circle signals in SLFN11-inducible cells in the absence or presence of doxycycline (Dox) induction using ImageJ. Data were obtained from two independent biological replicates, each with at least two technical replicates. Data are presented as mean ± SDs. ** indicates *p* < 0.01 (two tailed pair t-test).

C-circles are extrachromosomal, partially single-stranded telomeric DNA species that are highly enriched in ALT-positive cells and are widely used as a marker of ALT activity. To further determine whether SLFN11 regulates ALT activity, we examined C-circle levels in cells in the absence or presence of doxycycline (Dox)-induced SLFN11 expression using the C-circle assay.Consistent with active ALT, robust C-circle signals were detected in cells without SLFN11 induction. In contrast, induction of SLFN11 expression resulted in a reduction in C-circle levels (Figures 3C and D).

Together, these findings demonstrate that SLFN11 reduces both telomeric TERRA foci and extrachromosomal C-circle DNA, providing further evidence that SLFN11 acts as a negative regulator of ALT activity.

### SLFN11 causes telomere instability and reduces telomere DNA synthesis in ALT cells

ALT telomeres are characterized by persistent DNA damage signaling, including enrichment of γH2AX at telomeres under basal conditions ^26^. Previous studies have attributed process to replication impediments and formation of telomeric secondary structures such as G4 DNA and R-loops, which stall replication forks and promote DNA breaks ^27,57^. Despite this, ALT cells tolerate such stress through efficient DNA repair/recombination pathways. To explore this process, we stained U2OS expressing SLFN11 for telomeric γH2AX and 53BP1 foci. Expression of SLFN11 in U2OS cells markedly reduced γH2AX and 53BP1 accumulation at telomeres (Figures 4A, B, S3A and S3B). Since γH2AX and 53BP1 at telomeres can decrease under two non-mutually exclusive scenarios: (i) reduced telomeric DNA damage, or (ii) suppressed DNA damage signaling, and given the established role of SLFN11 as an inhibitor of DNA damage signaling and homologous recombination (HR) ^33,41,46^, we interpret the reduction in telomeric γH2AX upon SLFN11 expression as reflecting an attenuation of telomeric DDR signaling.

**Figure 4:**
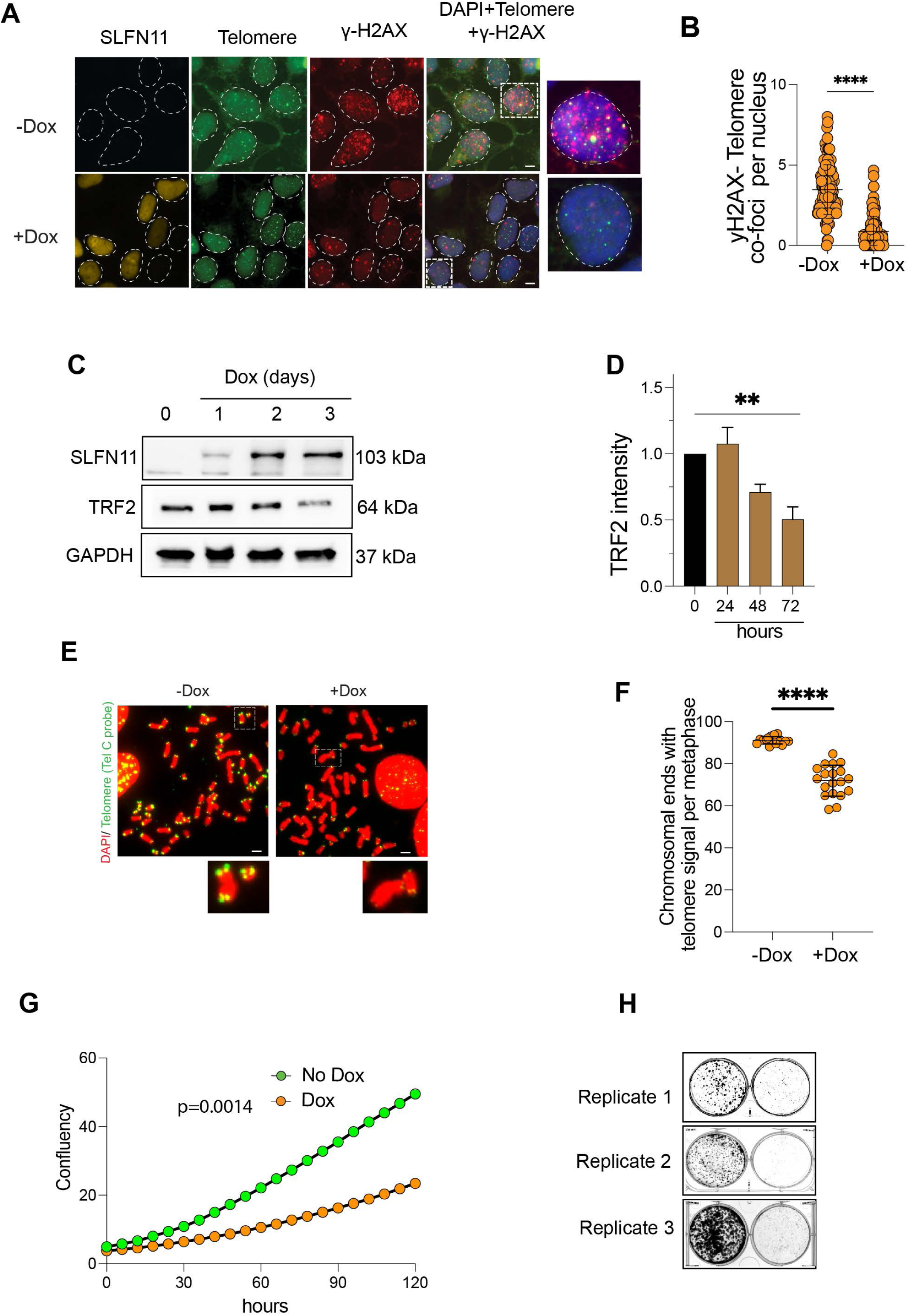
SLFN11 promotes telomere instability. **A.** Representative immunofluorescence combined DNA-FISH images of U2OS cells showing SLFN11-dependent suppression of γ-H2AX localization at telomeres after 4 days of doxycycline treatment. Scale bar, 5μm. **B.** Quantification of γ-H2AX foci colocalized with telomeres in U2OS cells without and with SLFN11 expression after 4 days of doxycycline treatment. At least 100 nuclei were counted for each condition in three independent biological replicates experiments. Data are presented as mean ± SD. ** indicates *p* < 0.01 (two tailed pair t-test). **C.** Representative Western blot showing reduced TRF2 protein levels following SLFN11 induction in U2OS cells. **D.** Quantification of TRF2 protein levels in SLFN11-inducible cells in the absence or presence of doxycycline (Dox) induction using ImageJ. Data were obtained from three independent biological replicates and are presented as mean ± SD. **p < 0.01 (two-tailed paired Student’s *t*-test). **E.** SLFN11 expression alters ALT telomeres examined by metaphase-FISH. Telomeres were stained using the TelC-Cy3 (F1002) telomere probe. DAPI was used to stain chromosomes (blue). Scale bar 5 μm **F.** Quantification of telomeric signals at chromosome ends in metaphase spreads of SLFN11-inducible cells in the absence or presence of doxycycline (Dox) induction. Data were obtained from three independent biological replicates and are presented as mean ± SD. **** indicates *p* < 0.0001 (two tailed pair t-test). **G.** Suppression of cell growth upon SLFN11 expression in U2OS. Cells were seeded at ∼10% confluency and treated with control or 5 µg/mL doxycycline to induce SLFN11. Real-time proliferation was monitored using the IncuCyte. The experiment was performed in three independent biological replicates. **** indicates *p* < 0.0001 (two tailed pair t-test). **H.** Suppression of colony formation upon SLFN11 expression in U2OS cells. Equal numbers of cells were seeded in 6-well plates and treated with control medium or 5 µg/mL doxycycline to induce SLFN11. Colonies were allowed to grow for 10 days, then fixed with methanol, stained with crystal violet, and imaged (n=3).

To further investigate whether SLFN11-mediated telomere destabilization affects DNA synthesis at ALT telomeres, we examined telomeric DNA synthesis during the G2 phase of the cell cycle, when ALT cells undergo recombination-associated telomere replication. We monitored EdU incorporation at TRF2-marked telomeres in cells with or without doxycycline (Dox)-induced SLFN11 expression. Cells were synchronized with thymidine for 18 hours, released for 4 hours, and subsequently treated with the CDK1 inhibitor RO-3306 for 18 hours to arrest cells in late G2 phase. Cells were then pulsed with EdU for 3 hours before fixation and immunofluorescence analysis (Figure S3C).Fluorescence imaging revealed EdU incorporation at TRF2-positive telomeric foci in cells without SLFN11 induction (Figures S3D and E). In contrast, SLFN11 induction reduced the number of EdU foci colocalizing with TRF2 (Figures S3B and C), indicating suppression of DNA synthesis at ALT telomeres.

Since our immunofluorescence analysis revealed reduced TRF2 signals at telomeres following SLFN11 induction, we next examined whether SLFN11 affects TRF2 protein expression. Western blot analysis confirmed that SLFN11 induction reduced TRF2 protein levels, further supporting its role in telomere destabilization (Figures 4C and D). Consistent with this, metaphase spreads following SLFN11 induction revealed the reduced telomere signal (Figures 4E and F), possibly indicative of unresolved telomeric DNA breaks leading to loss of telomeres

We also measured the growth of U2OS cells upon SLFN11 expression and observed reduced proliferation and reduced colony formation (Figures 4G and H). Together, these findings indicate that SLFN11 compromises telomere stability in ALT cells leading to impaired cell proliferation.

### SLFN11 inactivation promotes ALT induced by ATRX suppression

DNA-protein crosslinks (DPCs) represent abnormal covalent attachments of proteins to DNA, which can generate replication stress and DNA double-strand breaks^58,59^. A recent study showed that excessive DPCs at telomeres can promote telomere breaks and trigger the ALT pathway in cells lacking ATRX ^31^. However, this study was conducted in HeLa cells, which do not express SLFN11, limiting the ability to assess the potential contribution of SLFN11 in this context.

To investigate whether SLFN11 can suppress the activation of the ALT pathway, we employed DU145 telomerase-positive (non-ALT cells), which display high endogenous levels of SLFN11 ^37^ (Figures 1B and 5A). Treatment of these cells with a combination of siRNA to suppress ATRX and the potent PARP inhibitor talazoparib (BMN673) ^60^ did not result in an accumulation of single-stranded telomeric C-rich DNA (ss-teloC), a hallmark of ALT pathway initiation (Figures 5B and C) By contrast, DU145 cells lacking SLFN11 exhibited a different response. The same treatment led to the accumulation of ss-teloC in a fraction of cells, indicative of ALT pathway initiation (Figures 5B and C)).

**Figure 5:**
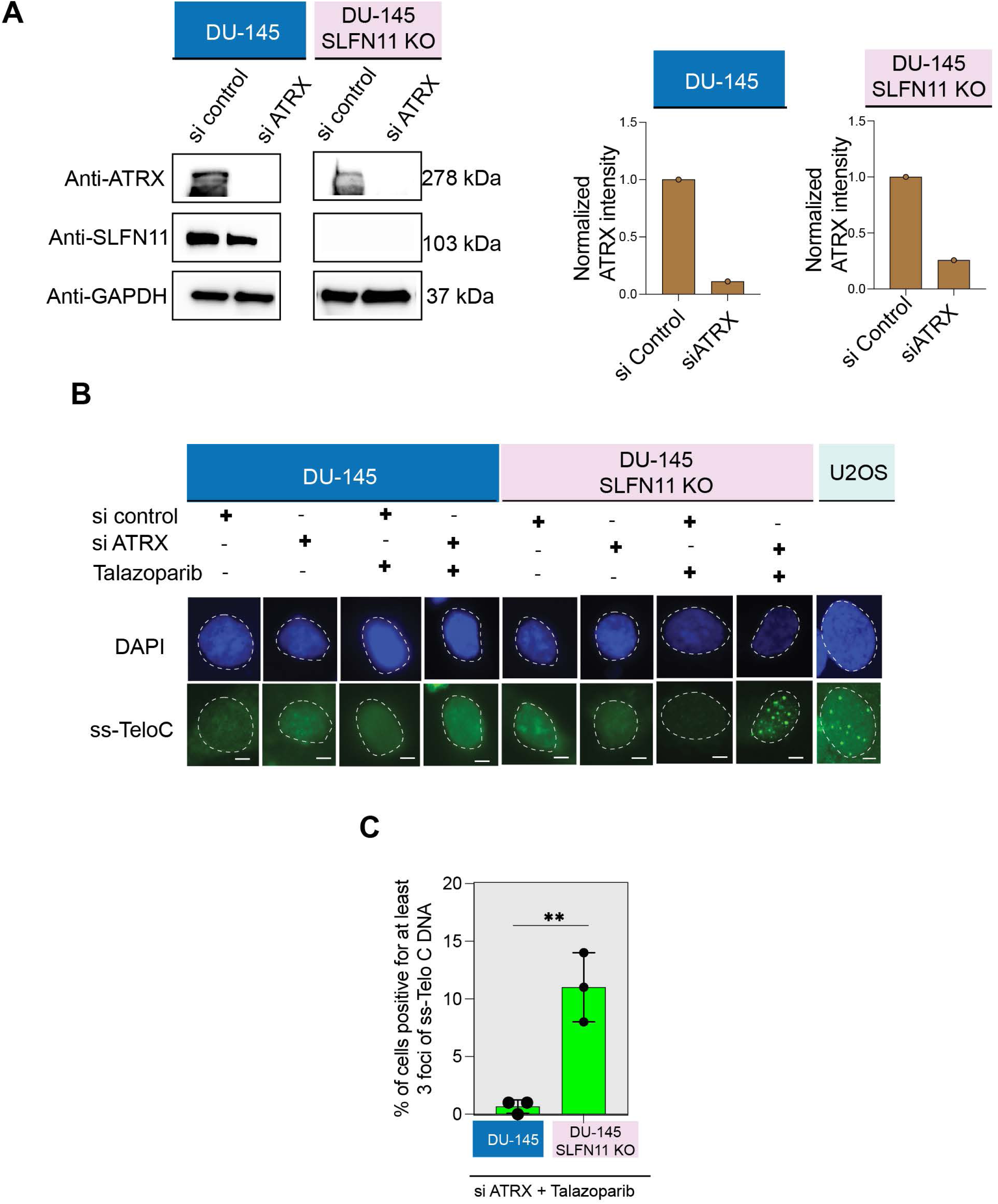
SLFN11 prevents ALT activation in ATRX-depleted DU145 cells treated with talazoparib. **A.** Representative Western blot demonstrating ATRX knockdown in DU145 SLFN11 wild-type and knockout cells. **B.** Representative immunofluorescence image showing single-stranded telomeric DNA (ssTeloC) signals detected by native FISH using a telomere-specific probe. DU145 WT and DU145 SLFN11 knockout cells were treated with siControl or siATRX for 72 hours and subsequently exposed or not to 5 µM talazoparib for 24 hours. The scale bar represents 5 µm. **C.** Quantification of ssTeloC foci in DU145 WT and DU145 SLFN11 knockout cells treated with or without siRNA control, siATRX and/or talazoparib At least 50 nuclei were counted for each condition in three independent experiments. ** indicates *p* < 0.01 (two tailed pair t-test).

To distinguish ALT activation from ss-teloC accumulation arising solely from increased telomeric replication stress, we examined the localization of ss-teloC relative to PML bodies. In SLFN11-KO DU145 cells, the induced ss-teloC foci colocalized with PML (Figures S4C) indicating the formation of ALT-associated PML bodies (APBs), a characteristic feature of ALT cells. These findings support the conclusion that the observed ss-teloC accumulation represents activation of the ALT pathway rather than merely increased telomeric replication stress.

We next performed a rescue experiment by generating SLFN11-KO DU145 cells carrying doxycycline-inducible SLFN11. Re-expression of SLFN11 markedly suppressed the appearance of ss-teloC foci following ATRX depletion and talazoparib treatment (Figures S4A, B and C)., demonstrating that the phenotype associated with SLFN11 loss could be reversed by restoration of SLFN11 expression.

To determine whether this effect was reproducible in an independent cellular background, we performed similar experiments in EW-8 cells. Consistent with our observations in DU145 cells, SLFN11-KO EW-8 cells exhibited ss-teloC foci that localized within PML, whereas this phenotype was not observed in their SLFN11-proficient counterparts (Figures S4D and S4E). Together, these results demonstrate that loss of SLFN11 facilitates ALT pathway activation under conditions of ATRX deficiency and replication-associated DNA damage, whereas SLFN11 expression suppresses this transition. These findings further support our observations in SLFN11-inducible U2OS cells and establish SLFN11 as a negative regulator of ALT.

## Discussion

The genetic reactivation of telomerase is a critical step for cancer cells to bypass senescence and achieve cellular immortality. Yet, a subset of cancers circumvents this process through an alternative telomerase-independent mechanism known as alternative lengthening of telomeres (ALT) ^9^. A central question is how ALT is activated or suppressed and what are the molecular factors that drive or oppose the transition of somatic cells into specialized ALT cancer cells with infinite replicative potential. Although most ALT tumors harboring inactivating mutations in ATRX or DAXX ^19,20,61,62^, genetic inactivation of ATRX alone cannot result in the induction or activation of ALT in telomerase-positive cancers or pluripotent stem cells ^31,63^, implying that additional molecular drivers control ALT.

In the present study, we identify SLFN11, an established determinant of sensitivity to replication damage-based chemotherapy ^37,46,64–67^ as being generally inactivated in ALT osteosarcoma cells and as a molecular suppressor of ALT. We provide evidence that SLFN11 suppresses ALT by binding to the telomeric regions of ALT cells. By generating a SLFN11 inducible doxycycline-system in ALT-positive U2OS cells, we investigated the localization and functions of SLFN11 at telomeres. SLFN11 induction was confirmed by immunoblotting and immunofluorescence (Figure 1). Although, distinct SLFN11 nuclear foci were observed in only a subset of ALT-positive cells, cells exhibiting SLFN11 foci consistently showed colocalization of these foci with telomeres (Figure 1D). Furthermore, proximity ligation assays (PLA) revealed a robust association of SLFN11 with the shelterin proteins TRF1 and TRF2 and RPA2 (Figure 1). The apparent difference between the frequency of detectable SLFN11-telomere colocalization by IF-FISH and the robust PLA signals may reflect differences in assay sensitivity. While IF-FISH requires sufficient local accumulation of SLFN11 to form detectable foci, PLA can detect close molecular proximity even at lower levels of SLFN11. Thus, SLFN11 may associate with telomeres more broadly than suggested by the frequency of detectable nuclear foci in IF-FISH experiments (Figure 1). Together, these findings provide evidence supporting the association of SLFN11 with telomeric chromatin in ALT cells. Similar to ATRX re-expression^25^, SLFN11 induction suppressed ALT activity, as evidenced by reductions in ALT-associated PML bodies (APBs), c-circles, and telomeric DNA synthesis (Figures 2, 3 and S3).

Because RPA maintains single-stranded telomeric DNA at sites of replication stress ^47^ and SLFN11 is known to bind RPA at stalled replication forks ^33,34^, we examined their interplay at ALT telomeres. SLFN11 showed colocalization with RPA and its re-expression not only disrupted RPA accumulation but also caused the disassembly of the BTRR (Bloom-TOP3A-RMI1-RMI2) complex and TRF2 destabilization, resulting in telomere deprotection (Figures 2 and 4).

Given that TERRA (Telomeric Repeat-containing Telomeric RNA) plays a central role in promoting ALT-associated recombination ^27,68,69^, we hypothesized that SLFN11 might regulate the ALT pathway by affecting TERRA. Consistent with this possibility, re-expression of SLFN11 significantly diminished TERRA RNA foci (Figure 3). Together, these findings demonstrate that SLFN11 suppresses ALT activity through impairment of TERRA R-loops and disassembly of the BTRR complex.

In our study, we provide evidence for a direct interference of SLFN11 with the telomeres of ALT cells. Loss of TRF2 is generally associated with telomere deprotection and the activation of telomere dysfunction-induced foci (TIFs). Surprisingly, despite observing reduced TRF2 levels following SLFN11 restoration, we did not detect an increase in TIF formation (Figure 4 and S3). Instead, we observed a reduction of TIFs, which normally accumulate in response to spontaneous telomere DNA damage in ALT cells due to stalled replication forks ^26^. One possible explanation is that SLFN11 attenuates DNA damage signaling pathways, including ATM- and ATR-mediated responses ^33,41,46^. Thus, even in the presence of reduced TRF2 levels, the downstream signaling required for robust TIF formation may be weakened when SLFN11 binds to the telomeres of ALT cells.

Metaphase chromosome spreads revealed a reduction of telomeric signals (Figures 4), indicating that SLFN11 expression induces telomere instability. A plausible model is that, in addition to suppressing the recruitment of homologous recombination factors to stalled replication forks ^33,46^, SLFN11 may also interfere with canonical DNA damage response (DDR) signaling at telomeres. Consequently, spontaneous DNA damage at telomeres may lead to telomeric DNA breaks without eliciting a corresponding increase in telomere dysfunction-induced foci (Figures 4 and S3). This could explain the apparent uncoupling of telomere structural instability from canonical DNA damage signaling in SLFN11-expressing cells.

Conversely, based on a recent publication ^31^ we established an experimental system to force ALT activation in telomerase-positive cells under defined ALT-permissive conditions. Specifically, ATRX-deficient telomerase-positive cells were exposed to talazoparib, which traps poly(ADPribose) polymerase complexes on DNA ^60^, blocks replication and induces DNA damage in a SLFN11-dependent manner ^70^. Under these conditions, ALT activation occurred exclusively in SLFN11-deficient cells, whereas SLFN11-proficient isogenic cell failed to initiate ALT activity (Figures 5 and S4). Collectively, these findings indicate that the combined loss of SLFN11 and ATRX establishes a permissive chromatin environment facilitating ALT pathway activation in the presence of telomeric DNA damage. A recent study supports our findings and conclusion that SLFN11 suppresses activation of the ALT pathway under DNA replication stress in ATRX-deficient cells by sensing ss-TeloC DNA ^71^.

Based on our results and the data provided simultaneously in a recent bioRxiv report ^71^, SLFN11 emerges as a novel identified suppressor of ALT by at least two non-exclusive mechanisms related to its binding to the telomeres of cells undergoing ALT. The first, which is described by the Boulton group in the bioRxiv publication ^71^ proposes that SLFN11 binding to ss-Telo C activates SLFN11’s endoribonuclease activity, leading to ribosomal stalling, global translational shutdown and apoptosis. The second, which we propose here in that SLFN11 directly disables the ALT molecular components including TRF2, APBs, TERRA and telomeric DDR, and suppresses the ALT pathway. We believe it is likely that both mechanisms are at play for SLFN11 to suppress the ALT-mediated proliferation of cancer cells.

Since SLFN11 is absent in nearly half of cancer cells, these new findings suggest that epigenetic inactivation of SLFN11 predisposes cancer cells to activate the ALT pathway upon exposure to telomeric DNA damage and conditions driving telomere decompaction such as ATRX deficiency. We surmise that restoration of SLFN11 expression in ALT cells, such as by reversing its epigenetic silencing may represent potent therapeutic strategy ^72,73^ to selectively target ALT positive tumors.

## Material and Methods

### Cell lines and cell culture conditions

The U2OS, DU145 and EW-8 cell lines were obtained from the Development therapeutics branch (NCI/NIH). The DU145 SLFN11 KO and EW-8 SLFN11 KO cell lines were previously generated and characterized in an earlier study from our laboratory^70^.U2OS cells were cultured and maintained in DMEM medium containing 10% Fetal Bovine Serum (FBS). DU145 and EW-8 cells were maintained in RPMI1640 medium supplemented with 10% FBS. During the study the cells were screened for mycoplasma using the MycoAlert Mycoplasma Detection Kit (LT07701 Lonza).

### Lentiviral transfection for stable cell lines

SLFN11 was cloned into pLVX-TetOne-Blast using In-Fusion Cloning System (Takara Bio) and primers (CCCTCGTAAAGAATTCATGGATTACAAGGATGACGAC and GAGGTGGTCTGGATCCCTAATGGCCACCCCACG according to manufacturer’s protocols. The lentiviral particles for the SLFN11 WT were generated using Lenti-X packaging single shots (Takara) and HEK293T cells and subsequently stored at-80° C. U2OS or DU145 SLFN11 KO cells were transduced with the lentiviral particles and selected with blasticidin for 10 days to generate the DOX-inducible SLFN11 cell line.

### Proximity ligation assays

Prior to proximity ligation assays (PLA), cells were fixed on coverslips using 4% paraformaldehyde in PBS for 10 minutes, then permeabilized with 0.2% TX-100 in PBS at room temperature. PLA was then performed using the Duolink^®^ PLA fluorescence protocol (Sigma). Briefly, fixed cells on coverslips were blocked for 1 hour at 37°C in the Duolink^®^ blocking solution (DUO82007). Primary antibodies were diluted in Duolink^®^ Antibody Diluent (DUO82008) and incubated 1 hour at room temperature. Coverslips were washed in 1x Wash Buffer A (DUO82049) twice for 5 minutes each, then incubated with Duolink^®^ In Situ PLA Probe Anti-Rabbit PLUS (DUO92002) and Anti-Mouse MINUS (DUO92004), diluted 1:5 in the Duolink^®^ Antibody Diluent, for 1 hour at 37°C. After washing samples in Wash Buffer A, ligation reactions were performed by diluting the Duolink^®^ Ligase 1:40 in 1x Duolink^®^ Ligation buffer and incubating for 30 minutes at 37°C. Following washing with Wash Buffer A, coverslips were incubated for 100 minutes at 37°C for the amplification reactions, which were performed by diluting the Duolink^®^ Polymerase 1:80 in 1x Duolink^®^ Amplification buffer using the FarRed detection kit (DUO92013). Samples were washed twice for 10 minutes each in 1x Wash Buffer B, once for 1 minute in 0.01x Wash Buffer B, then nuclei were stained with DAPI in PBS for 10 minutes at room temperature. Coverslips were mounted on slides using ProLong^TM^ Glass Antifade mountant (P36980) and cured for 24 hours prior to imaging. Images were captured using the ZEISS Axio Observer 7 by a blinded observer, and foci were counted using ImageJ.

### siRNA Knockdown

On-target plus siRNA smart pools targeting ATRX (*Dharmacon*, catalog no. L-006524-00-0005) were used for siRNA-mediated knockdown. Briefly, cells were seeded to achieve ∼30% confluency and transfected with siRNA using Lipofectamine RNAiMAX reagent (*Invitrogen*) according to the manufacturer’s instructions. Cells were incubated for the indicated time before being harvested for western blot or further processed for talazoparib treatment and/or IF-FISH.

### Immunofluorescence

Cells grown on glass coverslips were washed for 30 seconds with cyto buffer (100 mM NaCl, 300 mM sucrose, 3 mM MgCl₂, 10 mM PIPES, pH 6.8), followed by a 30 second wash in cyto buffer containing 0.5% Triton X-100, and again washed for 30 seconds in cyto buffer. Cells were then fixed in 4% paraformaldehyde (PFA) in PBS for 10 minutes at room temperature. After fixation, cells were washed with 0.5% PBST (PBS containing 0.5% triton-100) for 5 minutes and blocked for 30 minutes in BSA blocking buffer. Coverslips were incubated for 1 hour at room temperature with primary antibodies diluted in blocking buffer: TRF2 (*rabbit, 1:1000, NB110-57130, Novus Biologicals; mouse, 1:250, 4A794, Millipore Sigma*), PML (*mouse, 1:250, sc-966, Santa Cruz Biotechnology*), SLFN11 (*rabbit, 1:1000, 34858T, Cell signalling; mouse, 1:1000, sc-515071, Santa Cruz Biotechnology*), RPA2 (mouse, 1:250, sc-56770, *Santa Cruz Biotechnology*), BLM (*rabbit, 1:250, A-300-110, Bethyl Laboratories*), γ-H2AX (mouse, 1:500, 05-636, Sigma Millipore) and 53BP1 (1:500, NB 100-304, *Novus Biologicals)*. After washing with PBS (3×, 5 minutes each), coverslips were incubated with Alexa Fluor-conjugated secondary antibodies (*Alexa fluor 488, A11034; Alexa fluor 568, A11031; Thermofisher Scientific or Alexa fluor 647*) for 30 minutes at room temperature in blocking buffer. Further the cells were processed for DNA or RNA FISH or after washing with PBS, the cells were mounted using vectashield antifade medium containing DAPI (Vector Laboratories). Images were acquired using a ZEISS Axio Observer 7.

### Western blotting

Cells were cultured in a humidified incubator at 37 °C with 5% CO₂ until reaching approximately 70% confluency. Following siRNA transfection or doxycycline (5 μg/mL) induction for the indicated time intervals, cells were harvested, washed once with PBS, and lysed in NTEN buffer (20 mM Tris-HCl, pH 7.5, 300 mM NaCl, 1 mM EDTA, 0.5% NP-40) supplemented with protease inhibitor. Cell lysates were incubated for 1 hour at 4 °C with gentle rocking and clarified by centrifugation at 10,000 × g for 10 min at 4 °C. Protein concentrations were determined using the Bradford assay (Bio-Rad). Equal amounts of protein (30 μg) were resolved by SDS-PAGE on 4-20% Mini-PROTEAN TGX precast gels (Bio-Rad) and transferred onto PVDF membranes (Millipore). Membranes were blocked with 5% non-fat dry milk in PBST (PBS containing 0.1% Tween-20) for 1 hour at room temperature and incubated overnight at 4 °C with the indicated primary antibodies diluted in blocking buffer. After washing three times with PBST, membranes were incubated with HRP-conjugated secondary antibodies for 1 hour at room temperature. Immunoreactive bands were detected using enhanced chemiluminescence and visualized using a ChemiDoc imaging system (Bio-Rad). Band intensities were quantified using ImageJ software and normalized to GAPDH as indicated.

### IF-DNA FISH

Following immunofluorescence as described above, cells were hybridized at room temperature for 2 hours with an Alexa Fluor 488 labeled telomeric PNA probe (CCCTAA)₃ in hybridization buffer (70% formamide, 10 mM Tris-HCl, pH 7.5, 0.5% blocking solution) under denaturing conditions (72° C for 3 minutes). After hybridization, coverslips were washed twice for 15 minutes each in hybridization buffer. Nuclei were counterstained and mounted using vectashield antifade mounting medium containing DAPI (Vector Laboratories).

### IF-RNA FISH

Cells grown on glass coverslips were washed for 30 seconds with cyto buffer (100 mM NaCl, 300 mM sucrose, 3 mM MgCl₂, 10 mM PIPES, pH 6.8), followed by a 30 second wash in cyto buffer containing 0.5% Triton X-100, and again washed for 30 seconds in cyto buffer. Cells were then fixed in 4% paraformaldehyde (PFA) in PBS for 10 minutes at room temperature. After fixation, cells were washed with 0.5% PBST (PBS containing 0.5% triton-100) for 10 minutes. Immunofluorescence was performed using an anti-SLFN11 and TRF2 antibody as described above. Following antibody staining, coverslips were dehydrated sequentially in 70%, 90%, and 100% ethanol (2 minutes each), air-dried, and hybridized overnight at 37 °C with a Cy3-labeled telomeric PNA probe [TelC, (CCCTAA)₃] in hybridization buffer (2× SSC/50% formamide) under native condition. After hybridization, coverslips were washed sequentially three times for 5 minutes each in hybridization buffer, three times in 2× SSC, and once in 1× SSC at room temperature. Nuclei were counterstained and mounted using vectashield antifade mounting medium containing DAPI (Vector Laboratories).

### Metaphase spreads

Telomeric fluorescence in situ hybridization (FISH) on metaphase spreads was performed as previously described^12^, with minor modifications. Briefly, cells were treated with 0.2 µg/mL colcemid for 12 hours following to arrest cells in metaphase after 4 days of doxycycline induction. Cells were then harvested by trypsinization, washed once with PBS, and incubated in 75 mM KCl for 15-20 minutes at 37 °C to induce hypotonic swelling. After swelling, cells were fixed in methanol: acetic acid (3:1, v/v) with three sequential changes of fixative, and metaphase spreads were prepared by dropping the suspension onto clean glass slides under humidified chamber. Slides were hybridized with an TelC-Cy3 labeled telomeric PNA probe (F1002, PNA Bio) under denaturing condition, 72 °C for 3 minutes. Nuclei were counterstained with DAPI, and images were acquired using a ZEISS Axio Observer 7 microscope.

### C-circle assay

C-circle assay was performed as previously^74^, with C-circle abundance was measured using a rolling-circle amplification assay followed by dot-blot detection. Genomic DNA was treated with HinfI and AluI (4 U/µg DNA each) together with RNase A (25 ng/µg DNA) for 2 h at 37 °C. Subsequently, 16 ng of the digested DNA was subjected to rolling-circle amplification using EquiPhi29™ DNA Polymerase (Thermo Fisher Scientific) for 3 h at 45 °C. Parallel reactions lacking phi29 polymerase were included as negative controls. Following amplification, the reaction products and corresponding minus-enzyme controls were transferred onto a positively charged nylon membrane (Roche) and crosslinked. Telomeric DNA was detected by hybridization with a 3′ DIG-labeled (CCCTAA)3 telomeric probe (IDT) at 15 ng/mL. Signals were visualized using the DIG Nucleic Acid Detection Kit (Roche) according to the manufacturer’s instructions. Dot-blot signal intensities were quantified using ImageJ

### Cell synchronization and EdU staining

Cell synchronization followed by EdU staining was performed as described previously with minor modifications^75^. Cells were seeded onto coverslips 24 hours prior to thymidine treatment. For doxycycline-inducible experiments, doxycycline was added at the time of seeding and was maintained continuously throughout the synchronization and labeling procedure. Cells were first treated with thymidine for 18 h, washed, and released into fresh medium for 4 hours. Cells were subsequently treated with RO-3306 for 18 hours to enrich the population at the G2/M transition. Following synchronization, cells were exposed to EdU for 3 hours to monitor DNA synthesis. Cells were then fixed, and incorporated EdU was detected by Click-iT chemistry according to the manufacturer’s protocol. After completion of the EdU detection reaction, immunofluorescence staining was carried out as described above with Anti-TRF2 antibody, followed by mounting and fluorescence imaging.

### DPC mediated ALT induction and ss-telo C staining

Cells were grown on a prior to transfection with si control or siATRX for 48 hours. Followed cells were placed on cover glass followed by treatment with 5uM talazoparib for 24 hours. The concentration of 5 µM talazoparib (BMN 673) was selected based on a previous study by Clynes and colleagues, in which the same concentration was used to induce PARP trapping in HeLa cells ^31^. The cells were fixed 4% paraformaldehyde and subjected to ss-Telo C staining as described previously ^12^. Briefly, cells were incubated with RNase A (500 µg/mL) in blocking solution (1 mg/mL BSA, 3% goat serum, 0.1% Triton X-100, 1 mM EDTA in PBS) for 1 hour at 37 °C, dehydrated through an ethanol series (70%, 90%, and 100%), and hybridized with Alex flour CY3-labelled TelG (TTAGGG)ₙ PNA probe in hybridization buffer (70% formamide, 1 mg/mL blocking reagent, 10 mM Tris-HCl, pH 7.2). After hybridization, coverslips were washed twice in 70% formamide/10 mM Tris-HCl and three times with PBS, then mounted using vectashield antifade medium with DAPI. Images were acquired on a ZEISS Axio Observer 7 microscope.

### Colony formation assay

Equal numbers of cells were seeded into 6-well plates and allowed to adhere overnight at 37°C in a 5% CO₂ incubator. The following day, the medium was replaced with either fresh control medium or medium containing 5 μg/mL doxycycline to induce SLFN11 expression. Cells were cultured for 10 days, with medium replenished every 3 days to maintain induction and support colony growth. Followed, the colonies were fixed with methanol and stained with 0.5% crystal violet. Plates were rinsed with water to remove excess stain, air-dried, and imaged using a ChemiDoc imaging system (Bio-Rad).

### Incucyte live-cell growth assay

Cell proliferation was monitored in real time using the Incucyte® live-cell imaging system (Sartorius). Cells were seeded into 96-well tissue culture-treated plates at approximately 10% confluency in complete growth medium and allowed to adhere overnight at 37°C in a 5% CO₂ incubator. The following day, medium was replaced with fresh control medium or medium containing 5 μg/mL doxycycline to induce SLFN11 expression. Plates were then placed into the Incucyte® system, and phase-contrast images were acquired at defined time intervals using a 10× objective. Image analysis was performed using Incucyte® software to quantify percent confluence over time. For each experimental condition, percent confluence values from three technical replicate wells were averaged within each biological replicate. The mean values from three independent biological replicates were then plotted over time to generate cell proliferation curves.

### Quantification and statistical analysis

PLA foci were quantified using ImageJ (Fiji). IF-FISH foci and co-localized signals were counted by randomly selecting nuclei from multiple fields. Statistical analyses were performed using GraphPad Prism. Data are presented as mean ± standard deviation (SD) from the number of independent experiments indicated in each figure legend. p-values were calculated using a student’s t-test. Statistical significance is denoted as follows: ****p < 0.0001, ***p < 0.001, **p < 0.01 and n.s., not significant. Key resources table

### Key resource table

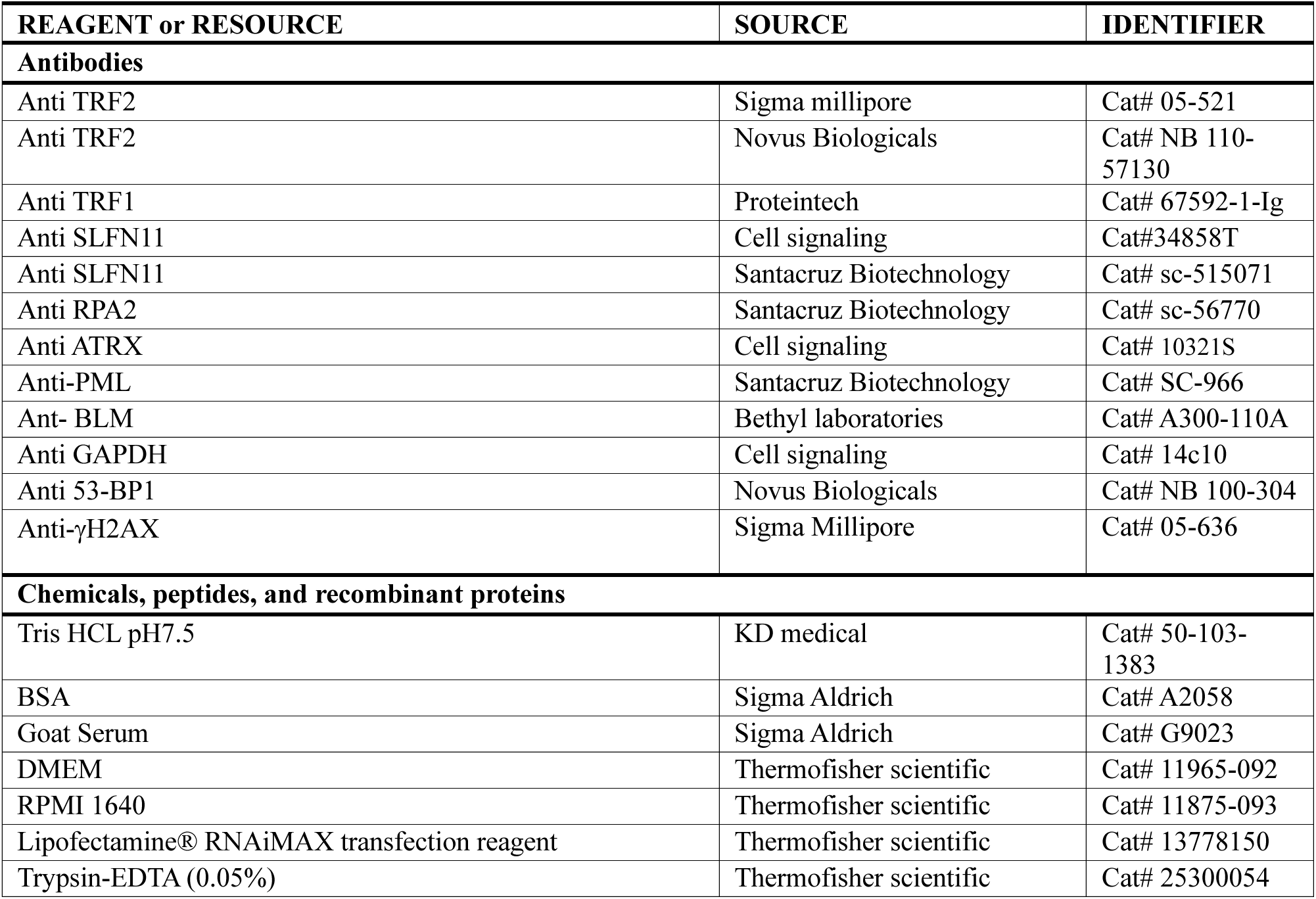
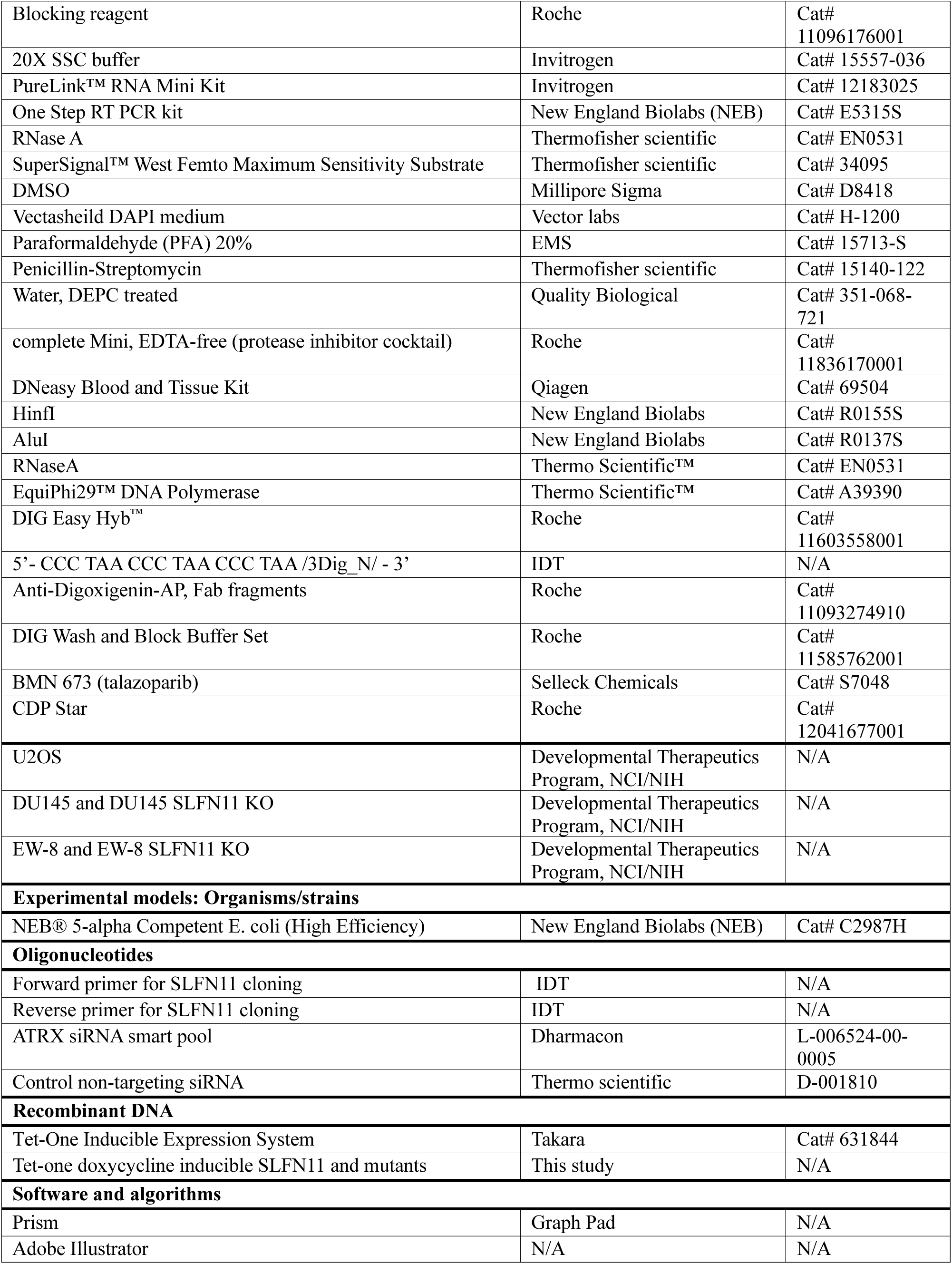
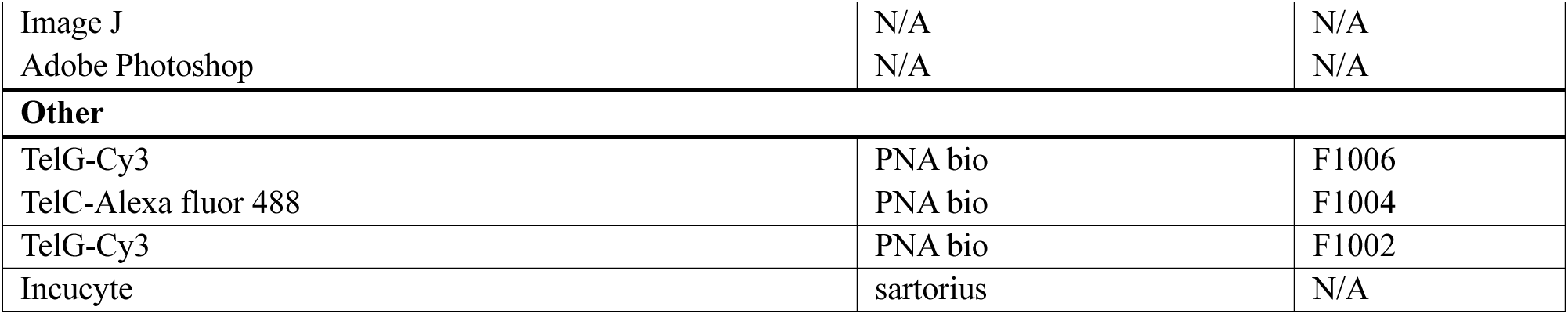

## Acknowledgements

This study was supported by the Center for Cancer Research, the Intramural Program of the National Cancer Institute, National Institutes of Health, Bethesda, Maryland 20892 (BC 006150). We acknowledge Dr. Mohit Arora for his suggestion on metaphase spread experiment and Dr. Haiqing Fu for telomere DNA synthesis experiment.

## Author contributions

P.K. and Y.P. designed research; P.K. performed research experiments, S.K. assisted in microscopy, A.L.W. and P.K. performed PLA assay, S.C and P. K. performed c-circle assay, P.K., S.Y.N.H. and D.T. generated doxycycline inducible SLFN11 cell lines; P.K., A.L.W., Y.M. and Y.P. analyzed data, P.K. and Y.P. wrote the manuscript.

---

**Supplementary Figure S1A:**
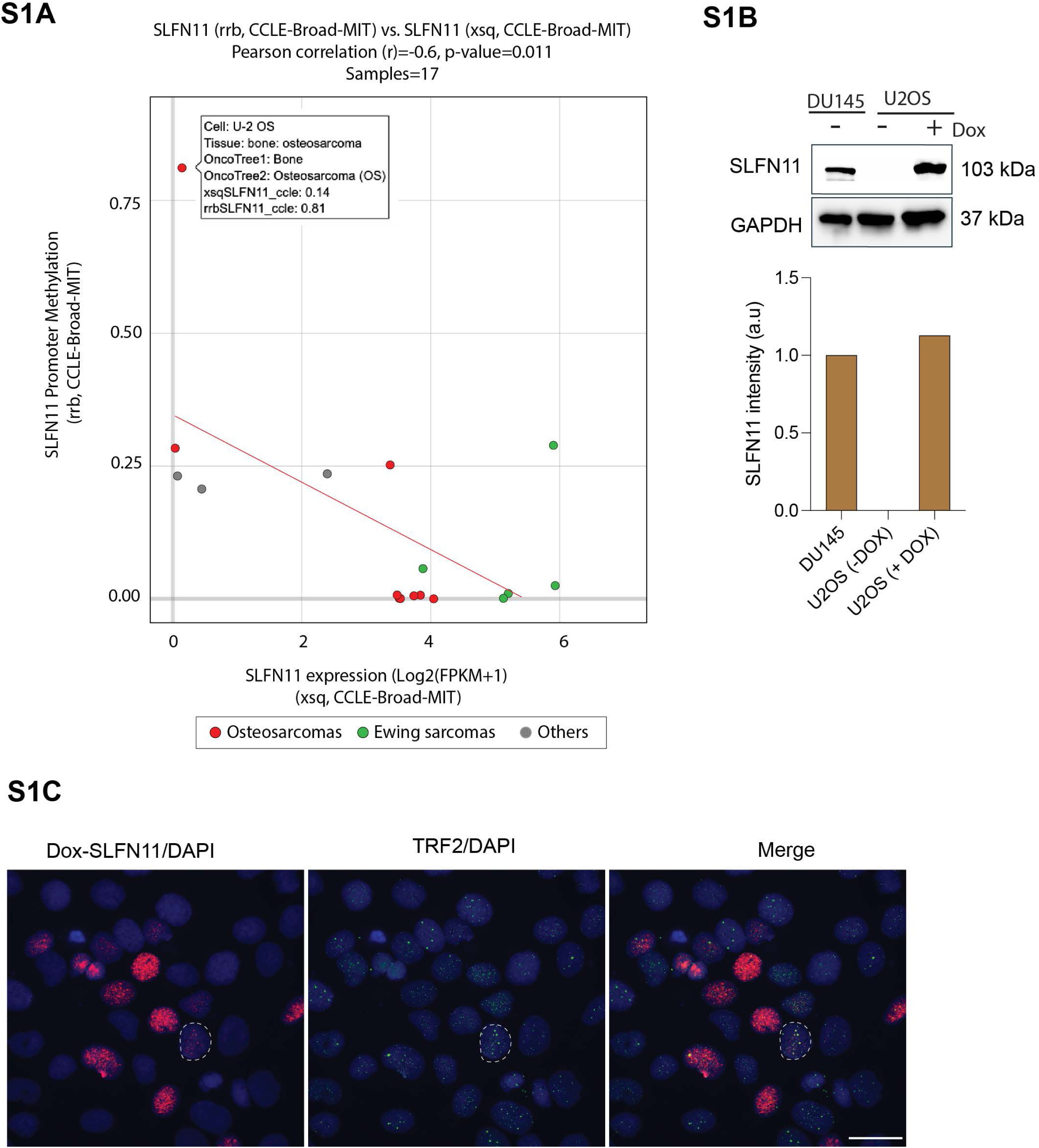
Methylation of *SLFN11* promoter in U2OS cells. Snapshot image of Sarcoma CellMinerCDB (https://discover.nci.nih.gov/sarcomaCellMinerCDB/) showing that high methylation of SLFN11 is correlated with low SLFN11 expression. **Supplementary Figure S1B:** Representative Western blot showing inducible expression of SLFN11 in U2OS cells treated with doxycycline (Dox) for 4 days, with DU145 cells included for comparison. **Supplementary Figure S1C. Localization of SLFN11 with TRF2.** A representative larger field of view showing that a small fraction of cells forms SLFN11 foci that colocalize with TRF2 following doxycycline treatment. The white-outlined cell is shown at higher magnification in Figure 1D. The scale bar represents 20 µm.

**Supplementary Figure S2.**
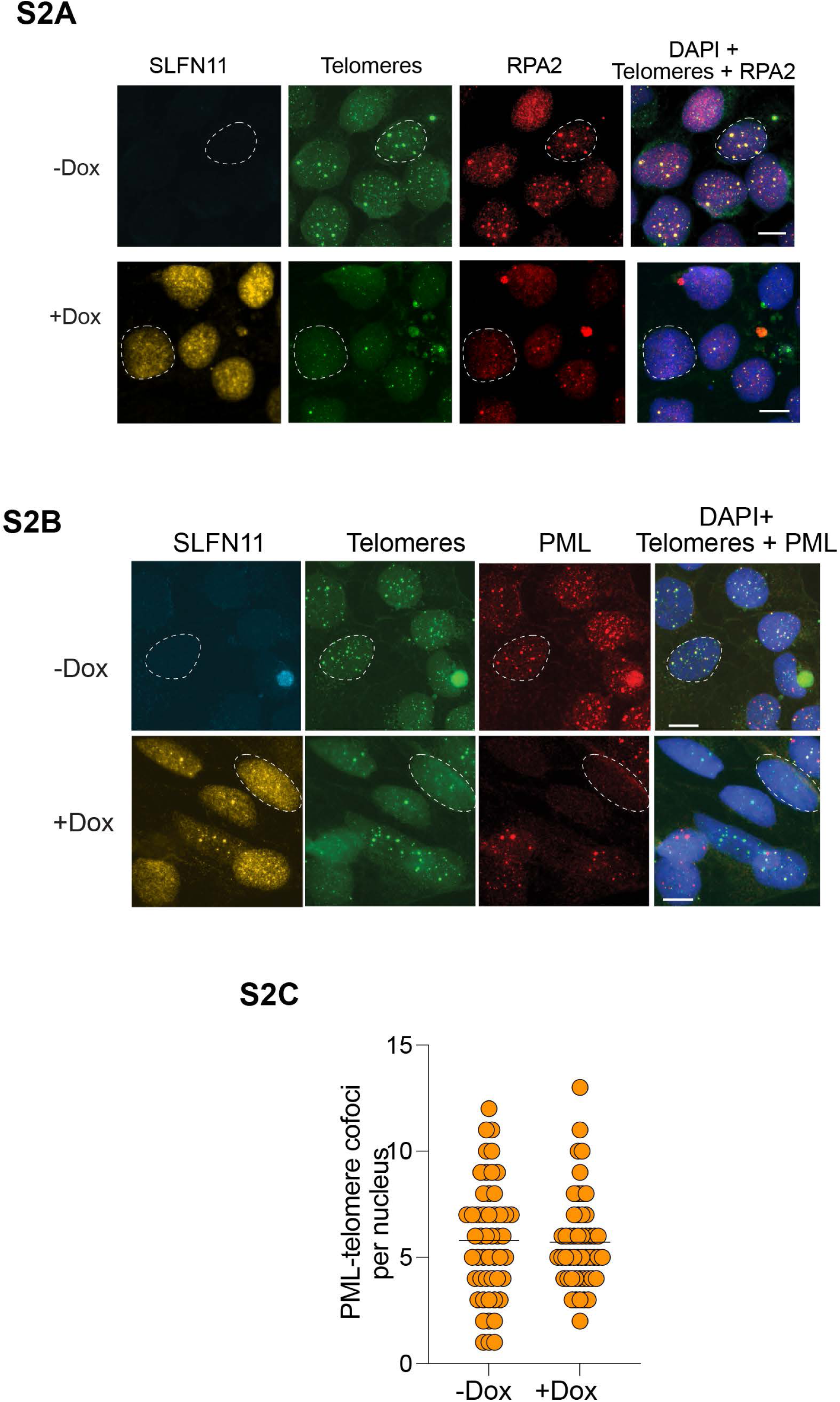
Localization of RPA2 and PML at telomeres. **A.** Representative large field views showing RPA2 localization at telomeres in the presence and absence of doxycycline treatment. The white-outlined cell is shown at higher magnification in Figure 2A. Scale bar, 20 µm. **B.** Representative large field views showing PML localization at telomeres in the presence and absence of doxycycline treatment. The white-outlined cell is shown at higher magnification in Figure 2E. Scale bar, 20 µm. **C.** Quantification of PML-Telomere co-foci in parental U2OS cells after doxycycline induction. At least 50 nuclei were counted for each condition from a single experiment.

**Supplementary Figure S3.**
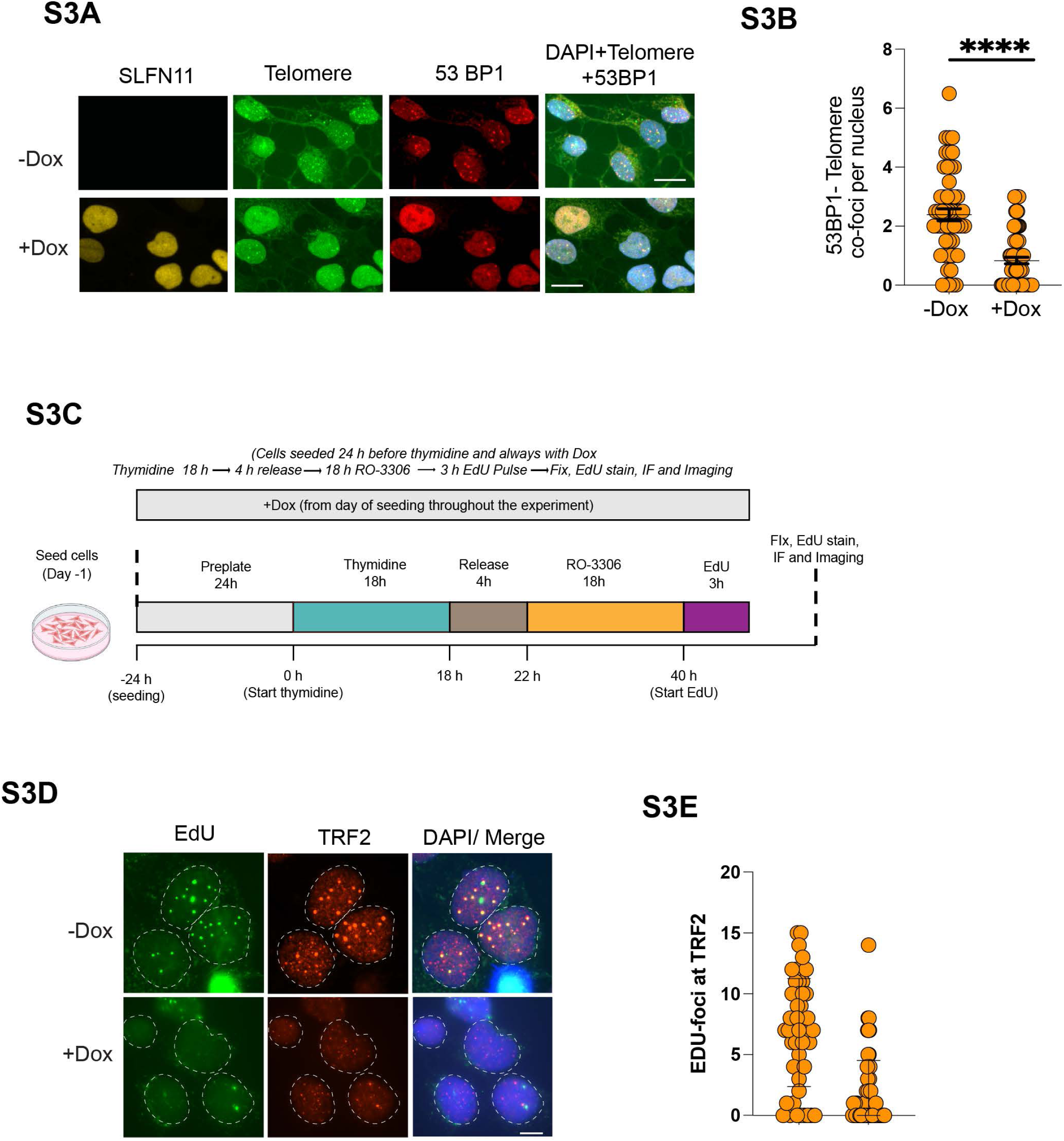
DNA damage response at telomere and telomere destabilization. **A.** Representative immunofluorescence combined DNA-FISH images of U2OS cells showing SLFN11-dependent suppression of 53BP1 localization at telomeres after 4 days of doxycycline treatment. Scale bar, 20 μm. **B.** Quantification of 53bp1 foci colocalized with telomeres in U2OS cells without and with SLFN11 expression after 4 days of doxycycline treatment. At least 50 nuclei were counted for each condition in two independent biological replicates experiments. **** indicates *p* < 0.0001 (two tailed pair t-test). **C.** Experimental scheme for analysis of telomeric DNA synthesis by EdU incorporation. Cells were seeded in the absence or presence of doxycycline 24 hours before thymidine treatment, with doxycycline maintained throughout the experiment. Cells were synchronized with thymidine for 18 hours, released into fresh medium for 4 hours and subsequently treated with the CDK1 inhibitor RO-3306 for 18 hours. Cells were then pulsed with EdU for 3 hours, followed by fixation, EdU staining, immunofluorescence (IF) for TRF2 and fluorescence imaging. **D.** Representative fluorescence images of U2OS cells without and with SLFN11 expression showing TRF2, EdU, and DAPI staining and merged images. Colocalization of EdU with TRF2 was used to identify DNA synthesis occurring at telomeres. **E.** Quantification of EdU foci colocalizing with TRF2 in −Dox and +Dox cells from one experiment (n=1). Each data point represents the number of EdU foci colocalized with TRF2 per nucleus. Data are from single experiment and are presented as mean ± SD.

**Supplementary Figure S4.**
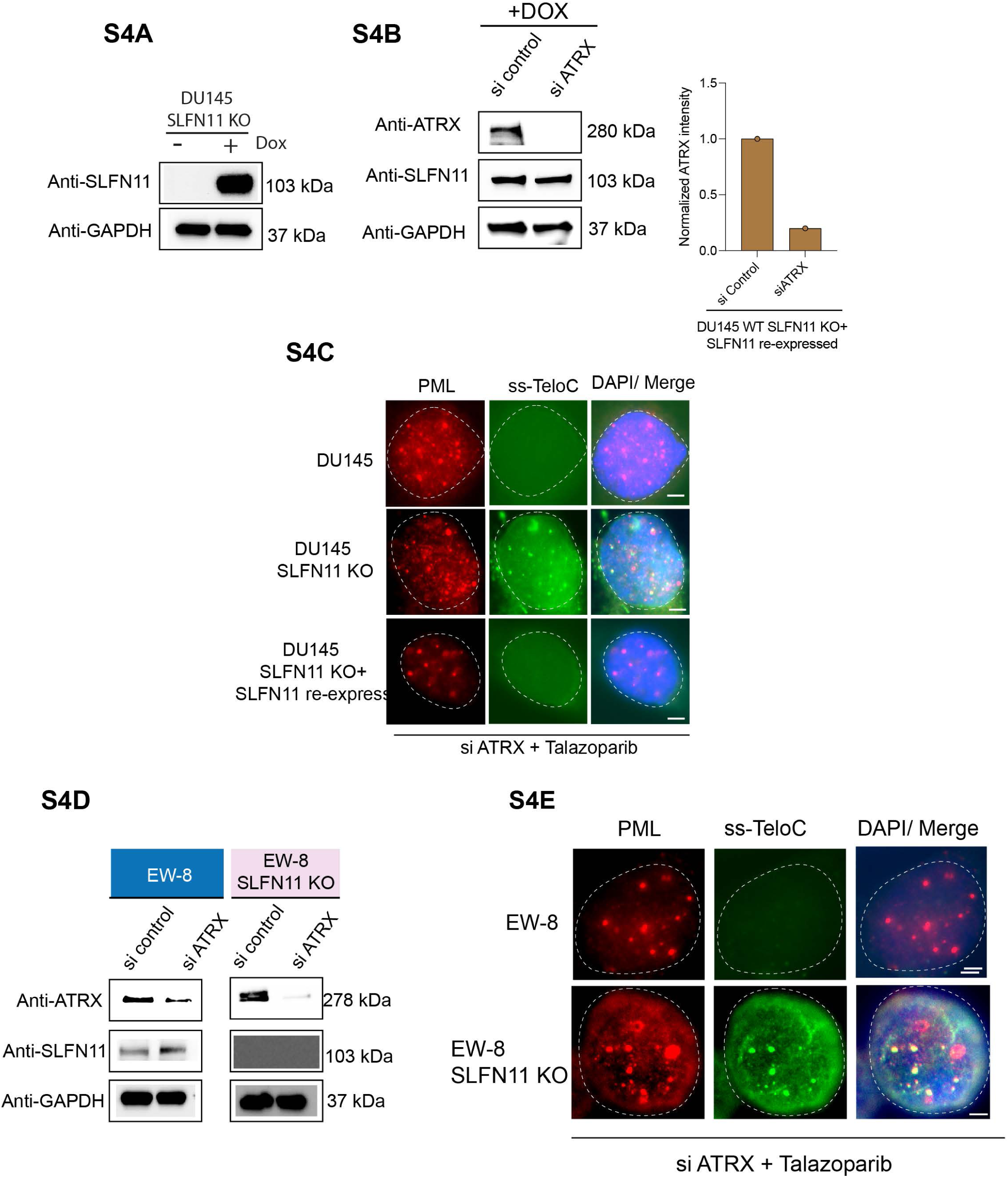
SLFN11 prevents ALT activation. **A.** Representative Western blot showing inducible expression of SLFN11 in DU145 SLFN11 KO cells treated with doxycycline (Dox) for 4 days. **B.** Representative Western blot demonstrating ATRX knockdown in DU145 SLFN11 KO with doxycycline inducible SLFN11 re-expression. **C.** Representative image showing single-stranded telomeric DNA (ssTeloC) signals detected by native FISH using a telomere-specific probe and PML using immunoflouresces in DU145 WT, DU145 SLFN11 knockout and DU145 SLFN11 KO with doxycycline inducible SLFN11 re-expression cells were treated with siControl or siATRX for 72 hours and subsequently exposed to 5 µM talazoparib for 24 hours. The scale bar represents 5 µm. **D.** Representative Western blot demonstrating ATRX knockdown in EW-8 SLFN11 wild-type and knockout cells. **E.** Representative image showing single-stranded telomeric DNA (ssTeloC) signals detected by native FISH using a telomere-specific probe and PML using immunoflouresces in EW-8 WT and EW-8 SLFN11 knockout cells were treated with siControl or siATRX for 72 hours and subsequently exposed to 5 µM talazoparib for 24 hours. The scale bar represents 5 µm.

